# FOXO1 inhibition and FADD knockdown have opposing effects on anticancer drug-induced cytotoxicity and p21 expression in osteosarcoma

**DOI:** 10.1101/2025.09.20.677161

**Authors:** Danielle Walker, Antanay Hall, Alexis Bonwell, Nancy Gordon, Danielle Robinson, Mario G. Hollomon

## Abstract

Forkhead box class O1 (FOXO1) and fas-associated death domain (FADD) regulate cell death pathways and homeostatic processes such as cell cycle progression and apoptosis. FADD phosphorylation promotes nuclear localization of FOXO1 and FOXO1 regulates FADD expression. Therefore, it is plausible that FOXO1 and FADD have synergistic or antagonistic effects on cell cycle regulation and the response to anticancer drug treatment in cancer cells. In the present study we report that AS1842856-mediated inhibition of FOXO1 reverses anticancer drug-induced cytotoxicity while FADD knockdown increases anticancer drug-induced cytotoxicity in osteosarcoma (OS). Reversed anticancer drug-induced cytotoxicity was accompanied with G2/M cell cycle arrest and increased expression of p21. The anti-cancer function of FOXO1 was further supported by the observation that OS cells that express higher basal levels of FOXO1 had increased sensitivity to camptothecin-induced cytotoxicity. FADD knockdown reversed FOXO1 inhibition-induced increase in p21 expression. The results presented in this study indicate that FOXO1 has a tumor suppressor function while FADD has a tumor promoting function in OS following anticancer drug treatment. The experimental approach used in this investigation also indicates that FADD antagonists the effect of FOXO1 on p21 expression in OS.

## Introduction

Osteosarcoma (OS) is the most prevalent bone cancer that primarily affects adolescents. While the overall survival rates for OS patients have improved over the past 30 years, improvement is still needed. To develop novel therapeutic options or improve current therapeutic options, a better overall understanding of OS biology is required. Forkhead box O1 is a transcription factor that regulates the expression of proteins involved in cell growth, differentiation, metabolism, apoptosis and redox homeostasis [1]. FOXO1 is expressed in bone and studies suggest that FOXO1 has a significant role in bone homeostasis and health [2]. For example, overexpression of FOXO1 has been reported to increase differentiation, proliferation, and migration of osteoblasts [3]. In addition, Teixeira et al. demonstrated that FOXO1 is a regulator of mesenchymal cell differentiation into osteoblasts [4]. Considering these critical functions in bone biology, it is plausible that FOXO1 has a major function in OS cell cycle regulation and response to anticancer drug treatment.

FADD is an adaptor protein that was first identified for its role in cell death, specifically apoptosis. FADD is also involved in other cell death pathways such as necroptosis [5] and autophagic cell death [6]. FADD has since been linked to non-death cellular processes such as cell proliferation [7], cell cycle regulation [8], cell differentiation [9] and protection against anticancer drug treatment [10]. Cellular localization is considered the determining factor for FADD-induced cell death or cell survival, with cytoplasmic localization associated with cell death and nuclear localization associated with cell survival [11]. p21 is a cyclin-dependent kinase inhibitor (CDKI) that inhibits all cyclin-dependent kinases (CDK) [12]. FOXO1 regulates the expression of p21 [13] and functional loss of p21 has been implicated in cancer [14]. FADD has been reported to regulate the cell cycle by regulating NFκB-mediated regulation of cell cycle cyclins [15].

Several studies have reported links between FOXO1 and FADD. For example, FADD is reported as a target gene of FOXO1 [17]) and FADD phosphorylation promotes FOXO1 nuclear translocation [17]. Considering the role of FOXO1 and FADD in cell death and cell cycle regulation and the reported associations between FOXO1 and FADD, it is plausible that FOXO1 and FADD have a linked effect on OS biology. The synergistic or antagonistic effect of FOXO1 and FADD on OS biology or the response of OS to anticancer drug treatment has not been explored.

The present study set out to investigate the effect of FOXO1 inhibition on cell cycle regulation and response to anticancer drug treatment in OS. The present study also investigated the combined effect of FOXO1 inhibition and FADD knock down on cell cycle regulation and anticancer drug-induced cytotoxicity. Here, we report that the inhibition of FOXO1 in OS decreases anticancer drug-induced cytotoxicity and induces G2/M cell cycle arrest. We also report that FOXO1 inhibition-mediated reversal in anticancer drug-induced cytotoxicity is cancer type-dependent. In addition, we present data that indicates that FOXO1 and FADD have opposing functions in cell cycle regulation and on the response to anticancer drug treatment in OS.

## 2. Results

### 2.1. OS cells express different basal levels of FOXO1

To begin the present study, basal levels of FOXO1 were determined. Western blot analysis revealed different basal levels of FOXO1 protein in OS. CCHOSD and HOS cells expressed high basal levels of FOXO1 protein (Figure 1A). Pancreatic cancer cell lines, AsPc1 and MiaPaCa2, also expressed different basal levels of FOXO1 protein. AsPc1 cells expressed high basal levels of FOXO1 protein while MiaPaCa2 expressed very low FOXO1 protein (Figure 1B).

**Figure 1.**
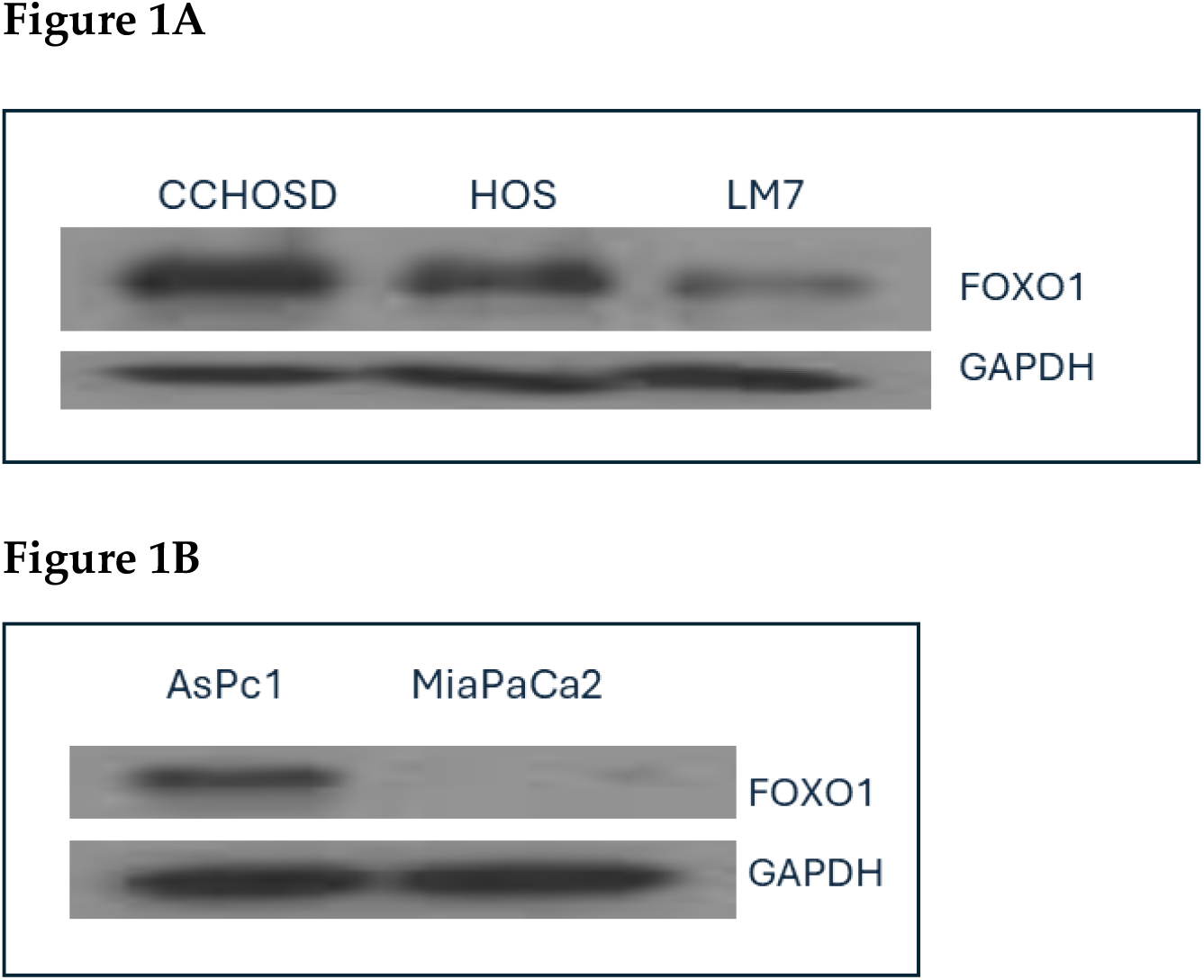
Osteosarcoma (OS) and pancreatic cancer cells express different basal levels of FOXO1 protein. Osteosarcoma (OS) cells were grown to approximately 70% confluency. Cells were next collected, lysed and total protein probed for FOXO1 protein expression. FOXO1 basal protein levels, **A**, OS cells, **B**, pancreatic cancer cells. GAPDH served as a protein loading control. Immunoblot is representative of immunoblots from two independent experiments.

### 2.2. FOXO1 inhibition induces selective cell death and increases p21 expression

AS1842856 is a specific inhibitor of FOXO1 that inhibits transcriptional activity by binding to the active, non-phosphorylated form of FOXO1 [18]. AS1842856 treatment induced higher overall cytotoxicity in pancreatic cancer cells compared to OS cancer cells (Figure 2A and 2B).

**Figure 2.**
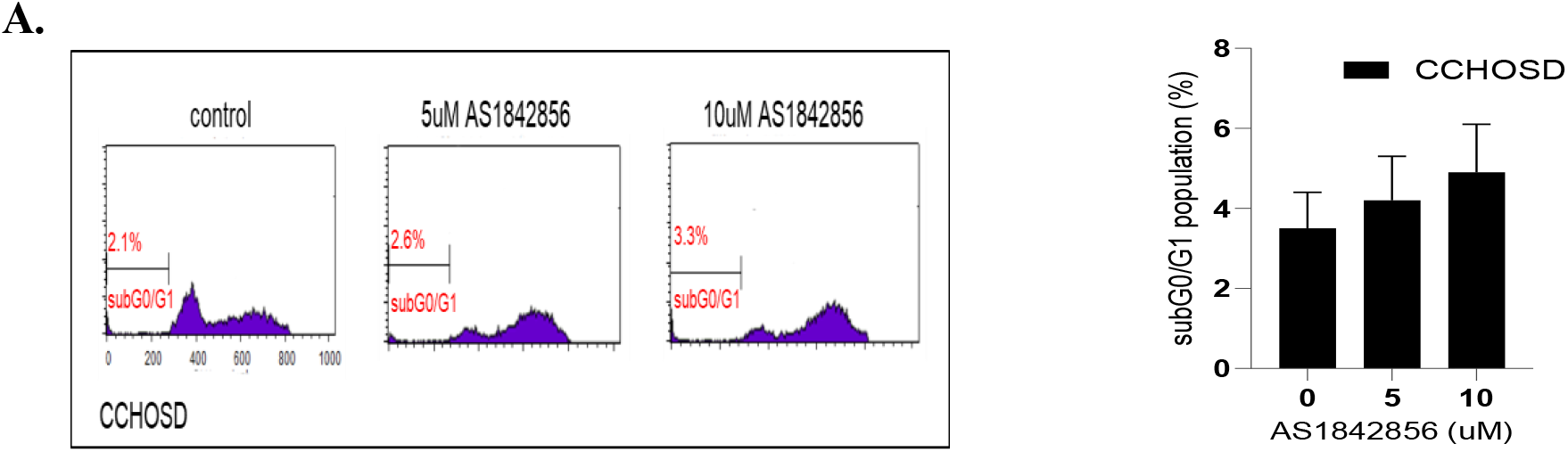
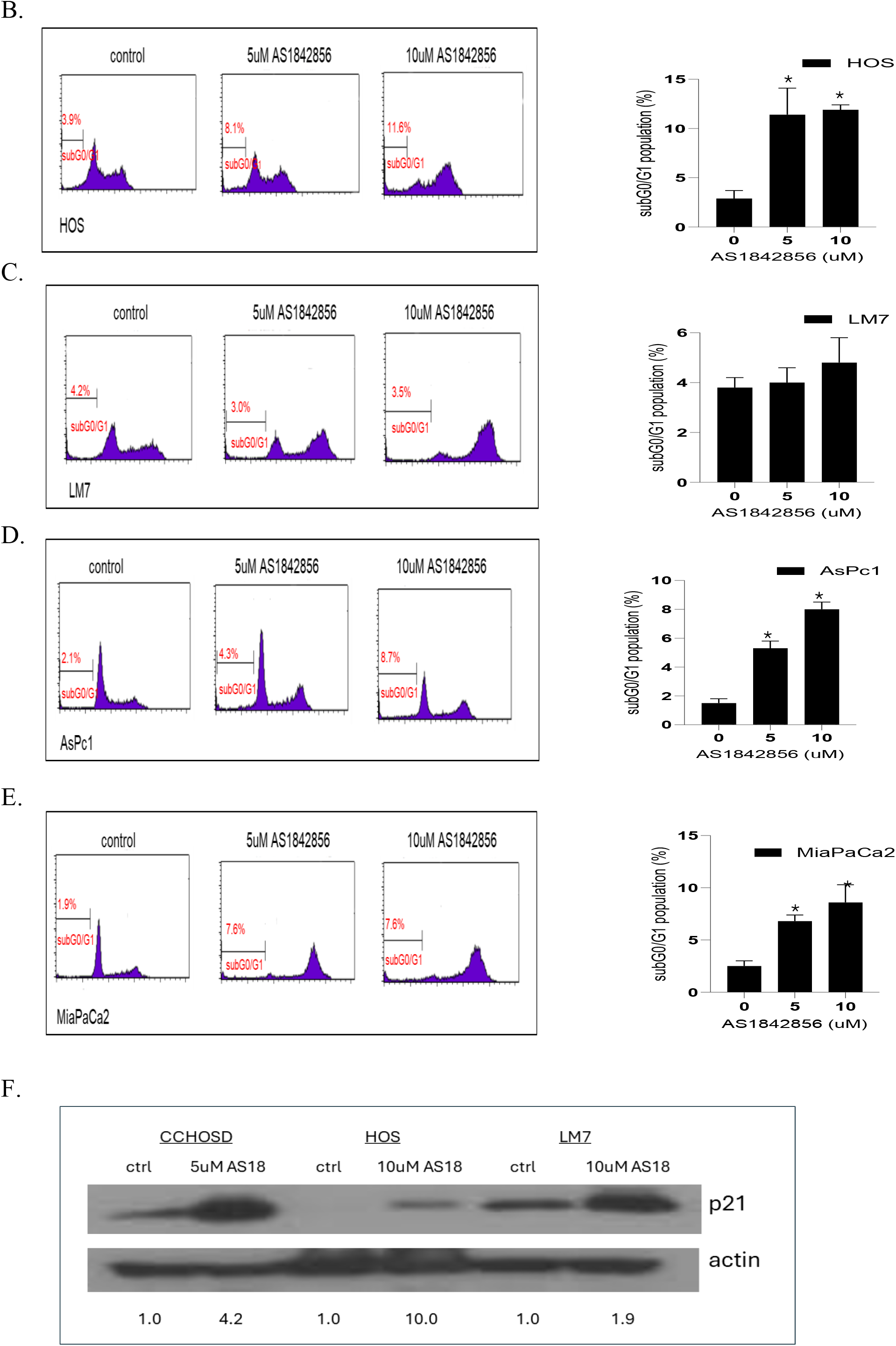
FOXO1 inhibition induces higher overall cytotoxicity in pancreatic cancer cells compared to OS cancer cells and increases p21 expression. Cells were treated with AS1842856 at concentrations indicated in figures. CCHOSD and HOS cells were treated with AS1842856 for 24 h. LM7, AsPc1, MiaPaCa2 cells were treated with AS1842856 for 48 h. Following AS1842856 treatment, cells were collected and processed for cell cycle analysis to determine the subG0/G1 cell population of cells. Results from one representative histogram is shown in the left panel. Mean results are shown in the right panel. **A**, CCHOSD, **B**, HOS, **C**, LM7; **D**, AsPc1, **E**, MiaPaCa2. **F.** AS1842856 treatment increases p21 expression. The expression of treatment group p21/actin ratio was determined by densitometry for each cell line and compared to control group p21/actin ratio which was normalized to the arbitrary value of one. Data represents the results of at least three independent experiments, + SEM. *, p<0.05 was considered significant.

### 2.3. Camptothecin induces significant cell death and increases p21 expression

CPT is a topoisomerase I inhibitor that binds to topoisomerase I and prevents DNA religation during DNA replication leading to DNA strand breaks and subsequent cell death [19]. CPT treatment induced significant cytotoxicity in all cell lines investigated (Figure 3).

**Figure 3.**
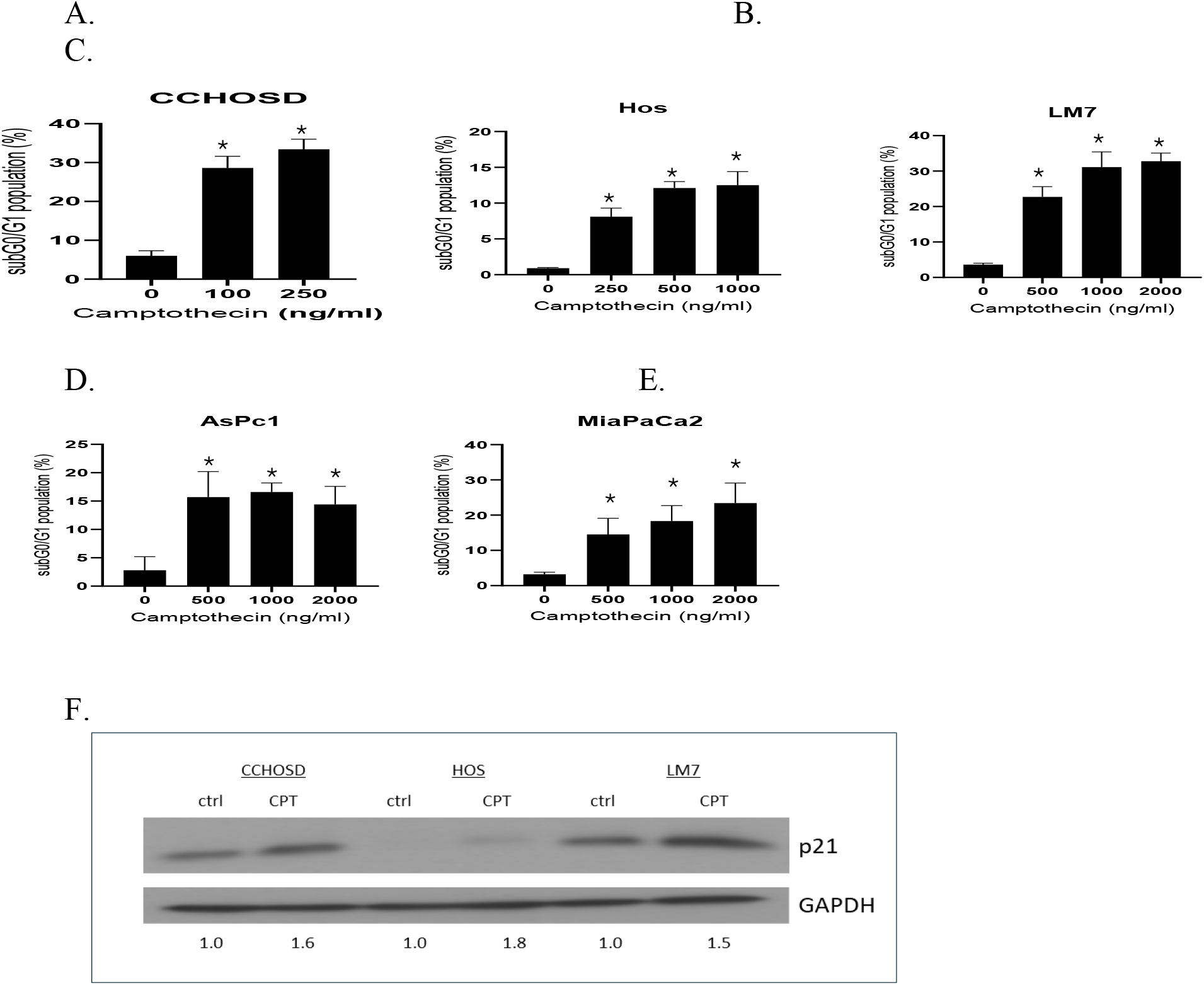
Camptothecin induces cytotoxicity and increases p21 expression. Cells were treated with CPT at concentrations indicated in figures. CCHOSD and HOS cells were treated with CPT for 24 h. LM7, AsPc1 and MiaPaCa2 cells were treated with CPT for 48 h. Following CPT treatment, cells were collected and processed for cell cycle analysis to determine the subG0/G1 cell population of cells. **A**, CCHOSD, **B**, HOS, **C**, LM7, **D**. AsPc1, **E**, MiaPaCa2. **F**. CPT treatment increases p21 expression. CCHOSD, 100ng/ml CPT; HOS, 500ng/ml CPT; LM7, 2000ng/ml. The expression of treatment group p21/actin ratio was determined by densitometry for each cell line and compared to control group p21/actin ratio which was normalized to the arbitrary value of one. Data represents the results of at least three independent experiments, + SEM. *, p<0.05 was considered significant.

### 2.4 FOXO1 inhibition reverses CPT-induced cytotoxicity and FADD knockdown increases CPT-induced cytotoxicity

To investigate the effect of FOXO1 inhibition on anticancer drug-induced cytotoxicity, OS cells were pretreated with AS1842856 followed by treatment with CPT. FOXO1 inhibition reversed CPT-induced cytotoxicity in all OS cell lines investigated (Figure 4). Conversely, FADD knock down increased OS sensitivity to CPT-induced cytotoxicity (Figure 5A). These results indicate that FADD has a protective role in OS following CPT treatment while FOXO1 has a tumor suppressor role following CPT treatment. FOXO1 inhibition reduced the level of FADD knockdown-induced cytotoxicity (Figure 5B).

**Figure 4.**
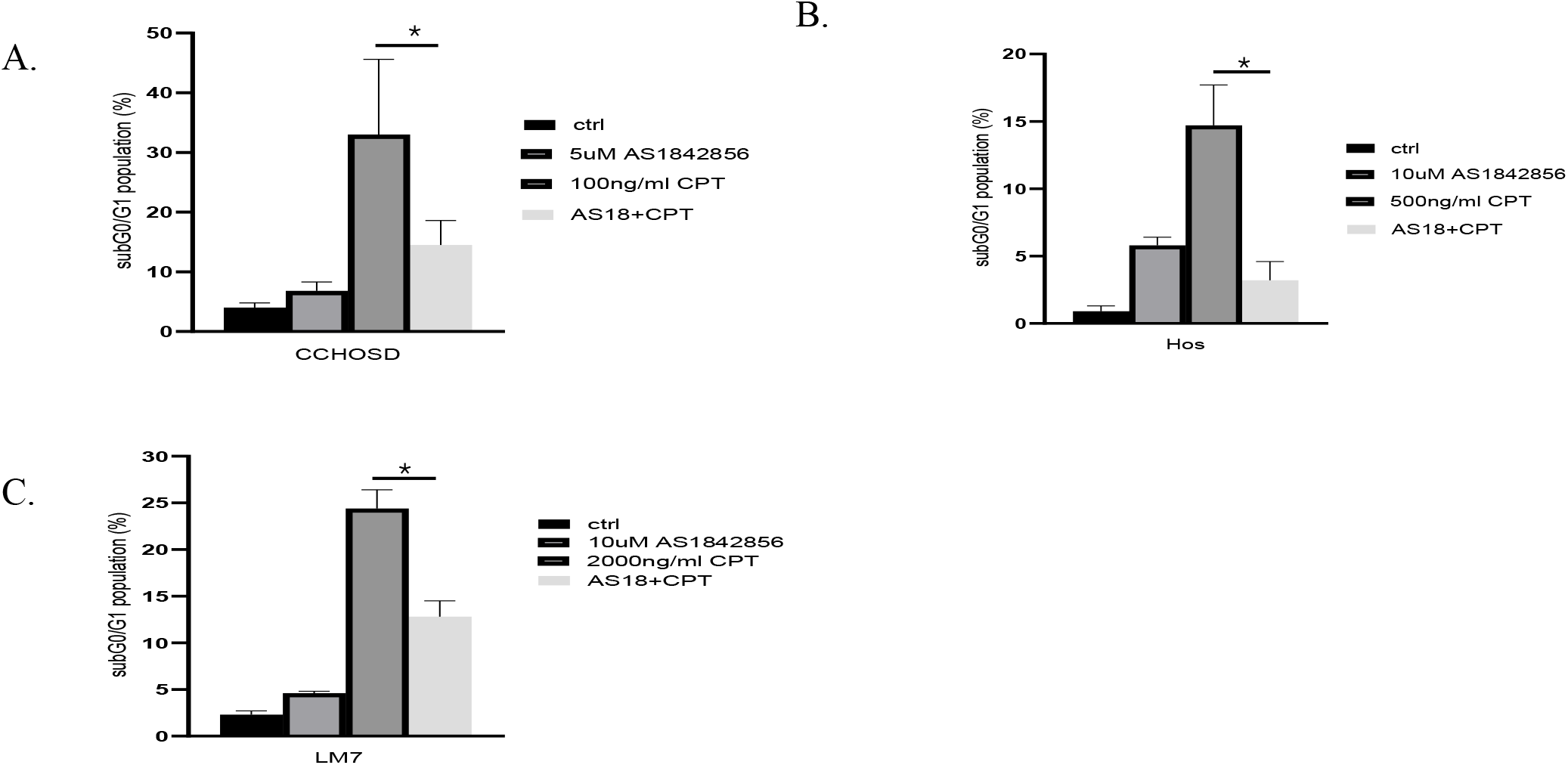
AS1842856 reverses CPT-induced cell cytotoxicity in OS cells. Cells were pretreated with AS1842856 followed by CPT treatment at concentrations indicated in figures. CCHOSD and HOS cells were treated with CPT for 24 h. LM7 cells were treated with CPT for 48 h. Following drug treatment, cells were collected and processed for cell cycle analysis to determine the subG0/G1 cell population of cells. **A**, CCHOSD, **B**, HOS. **C**, LM7. Data represents the results of at least three independent experiments, + SEM. *, p<0.05 was considered significant.

**Figure 5.**
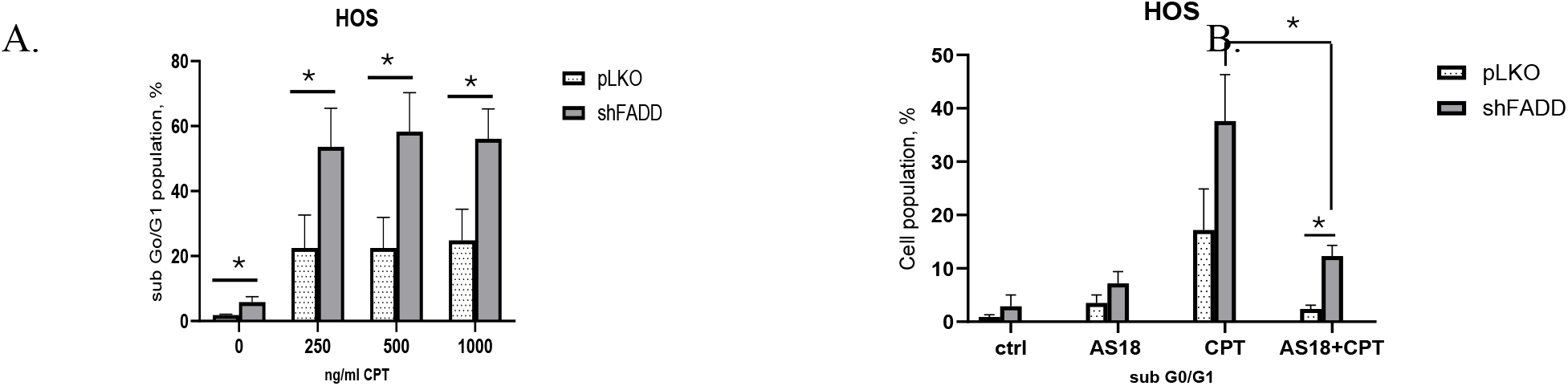
FADD knock down increases CPT-induced cytotoxicity which is reversed by FOXO1 knockdown. **A**, Wildtype (pLKO) or FADD knockdown (shFADD) cells were treated with CPT for 24h. **B**, Wildtype (pLKO) or FADD knockdown (shFADD) cells were pretreated with AS1842856 followed by CPT. Following drug treatment, cells were collected and processed for cell cycle analysis to determine the cell population in the subG0/G1phase. Data represents the results of at least two independent experiments, + SEM. *, p<0.05 was considered significant.

### 2.5 FOXO1 inhibition does not reverse CPT-induced cytotoxicity in pancreatic cancer cells

To determine if the FOXO1 inhibition-mediated reversal in CPT-induced cytotoxicity was restricted to OS, two pancreatic cancer cell lines were pretreated with AS1842856 followed by CPT treatment. FOXO1 inhibition did not reverse CPT-induced cytotoxicity in AsPc1 or MiaPaCa2 pancreatic cancer cells (Figure 6). This observation suggests that the effect of FOXO1 inhibition on anticancer drug-induced cytotoxicity is cancer type-dependent.

**Figure 6.**
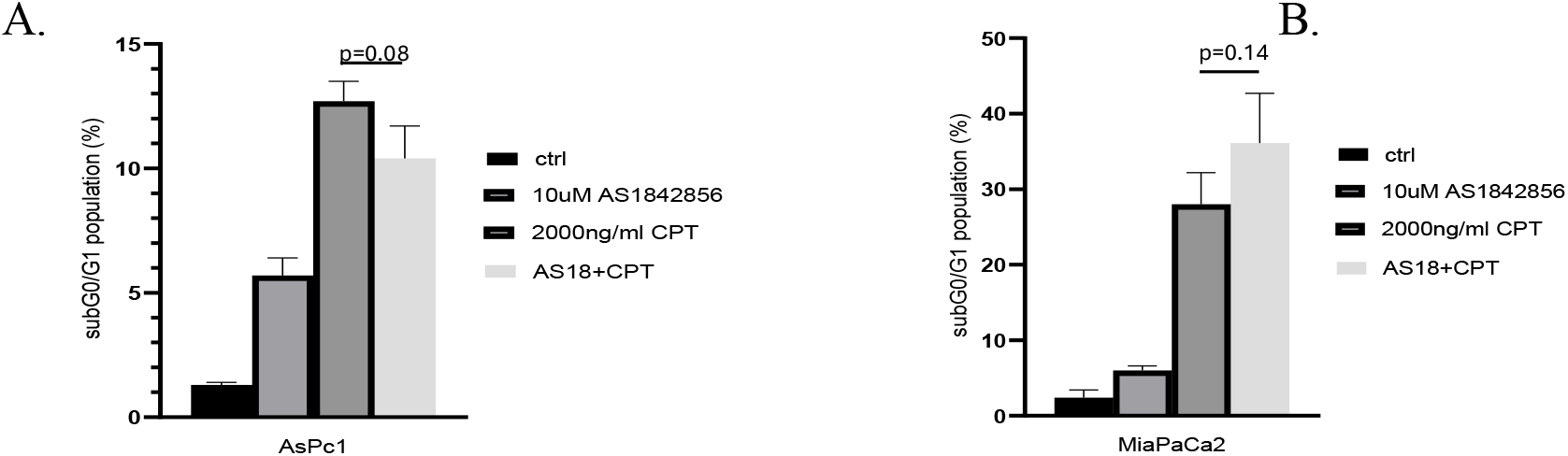
FOXO1 inhibition does not reverse CPT-induced cell death in pancreatic cancer cells. Cells were pretreated with AS1842856 followed by CPT treatment for 48 h. Following drug treatment, cells were collected and processed for cell cycle analysis to determine the subG0/G1 cell population of cells. **A**, AsPc1, **B**, MiaPaCa2. Data represents the results of at least three independent experiments, + SEM. *, p<0.05 was considered significant.

### 2.6 FOXO1 inhibition induces G2/M cell cycle arrest

To explore a possible link between FOXO1 inhibition-mediated reversal of CPT-induced cytotoxicity and cell cycle status, the effect of FOXO1 inhibition on cell cycle progression was investigated. FOXO1 inhibition induced G2/M cell cycle arrest in all OS and pancreatic cancer cell lines investigated (Figure 7).

**Figure 7.**
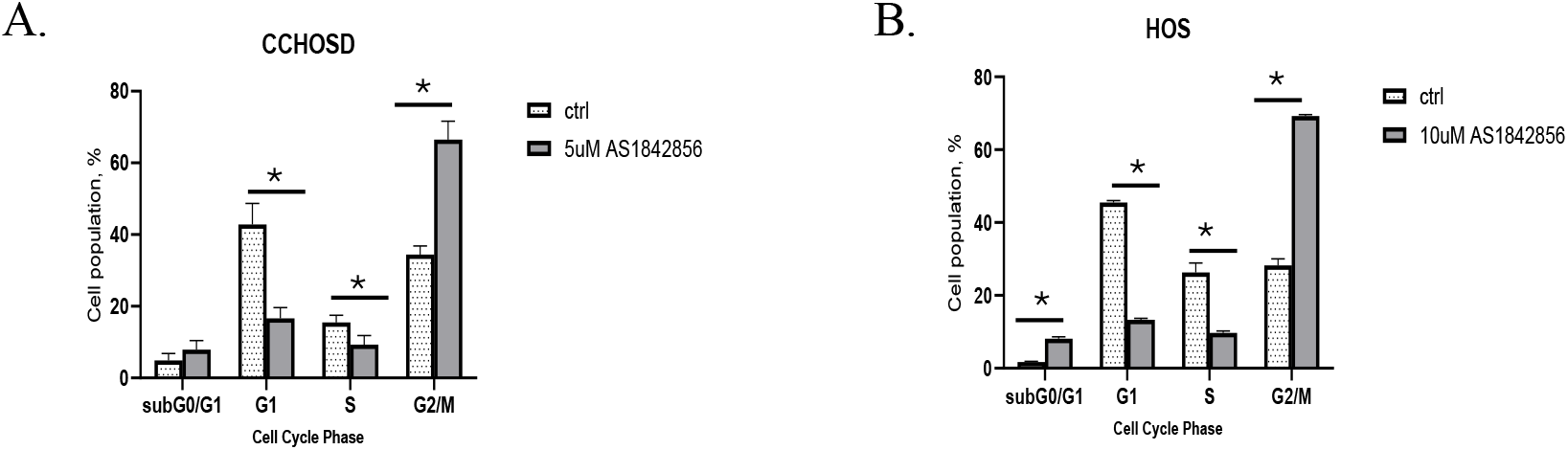
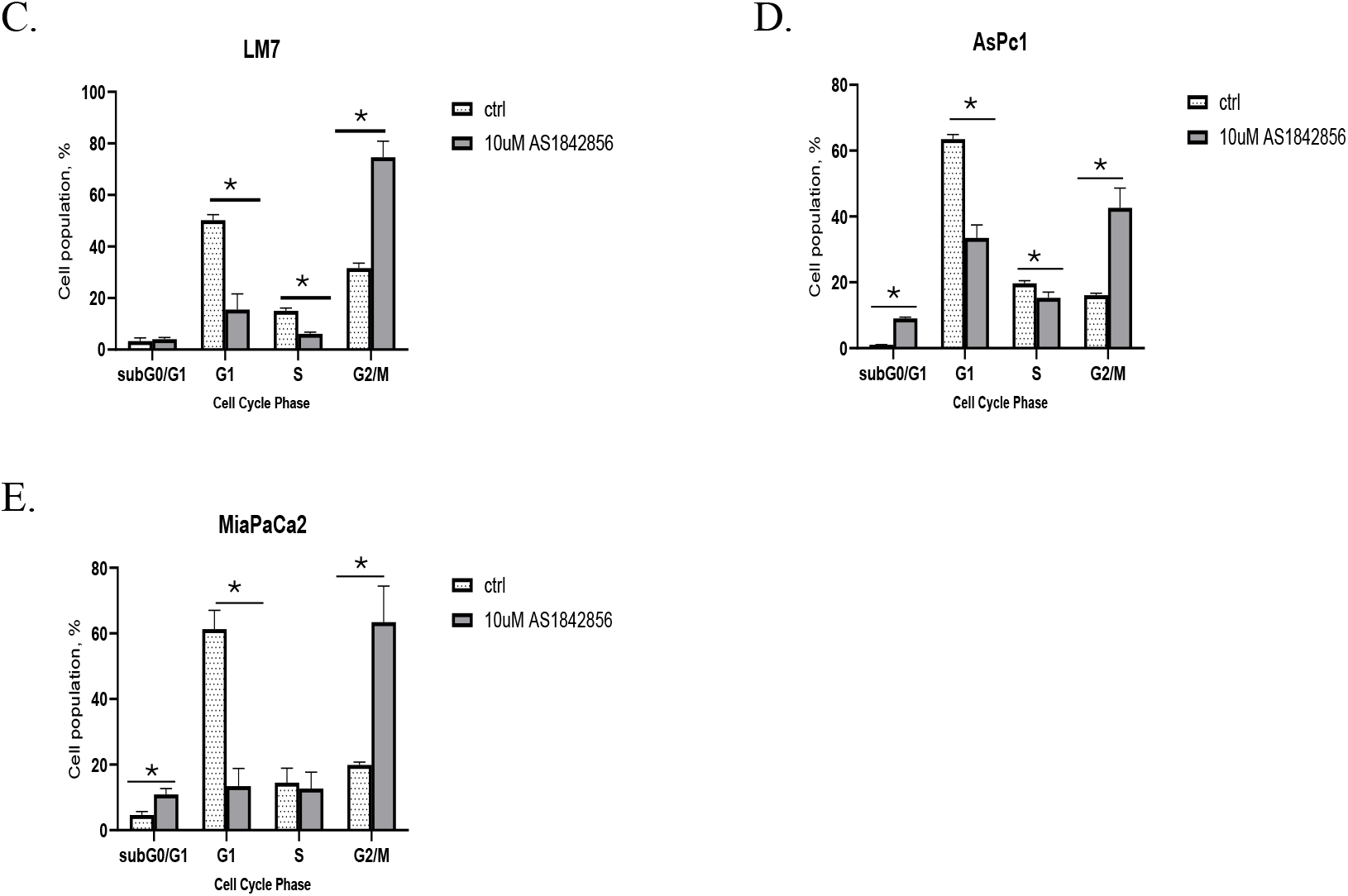
FOXO1 inhibition induces G2/M cell cycle arrest. CCHOSD and HOS cells were treated with AS1842856 for 24 h. LM7, AsPc1 and MiaPaCa2 cells were treated with AS1842856 for 48 h. Following AS1842856 treatment, cells were collected and processed for cell cycle analysis to determine the cell population in each cell cycle phase. **A**, CCHOSD, **B**, HOS. **C**, LM7., **D**, AsPc1, **E**, MiaPaCa2. Data represents the results of at least three independent experiments, + SEM. *, p<0.05 was considered significant.

### 2.7 FADD knock down alters p21 expression and reduces FOXO1 inhibition-induced p21 expression

To investigate the molecular mechanism for FOXO1 inhibition-mediated reversal in CPT-induced cytotoxicity and cell cycle arrest, basal protein expression of p21 was determined. CCHOSD and LM7 cells express higher basal levels of p21 compared to HOS (Figure 8). Western blot revealed a cell line-dependent effect of FADD knockdown on p21 expression in OS (Figure 9B). FADD knock down reduced p21 expression in untreated CCHOSD cells and increased p21 expression in untreated LM7 cells (Figure 9B). The effect of combined FOXO1 inhibition and FADD knock down on p21 expression was also determined. FADD knock down reversed the FOXO1 inhibition-induced increase p21 expression (Figure 9C-9E). These results suggest opposing roles of FOXO1 and FADD on p21 expression in OS.

**Figure 8.**
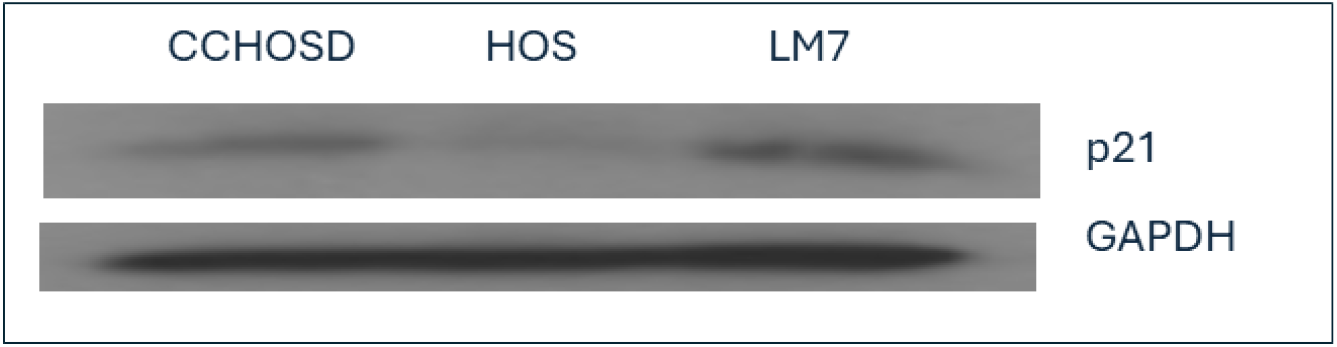
Osteosarcoma cells express different basal levels of cell cycle inhibitors. Osteosarcoma cells were grown to approximately 70% confluency. Cells were next collected, lysed and total protein probed for p21 protein expression. GAPDH served as a protein loading control. Immunoblot is representative of immunoblots from two independent experiments.

**Figure 9.**
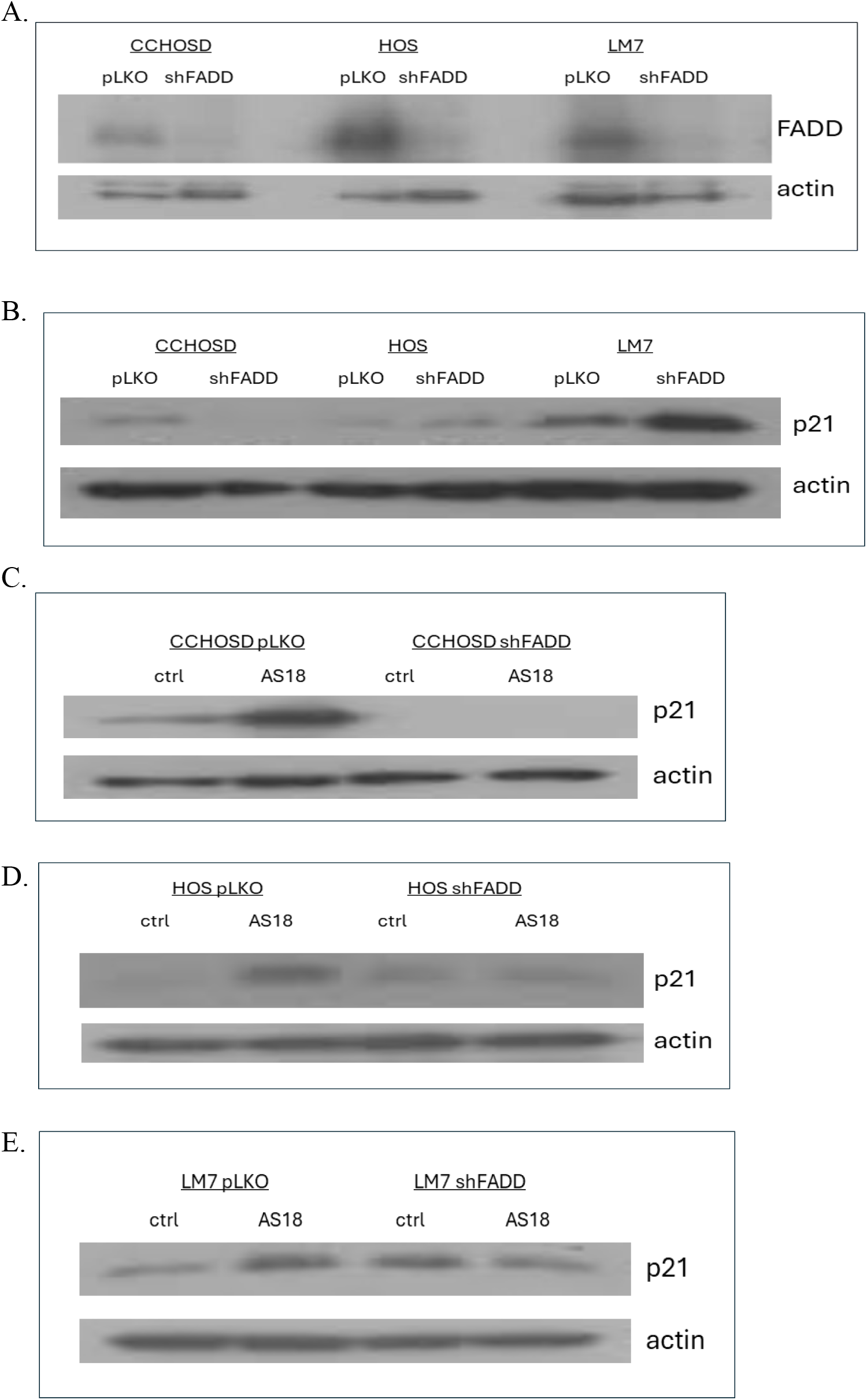
FADD knockdown reduces FOXO1 inhibition-induced p21 expression. Osteosarcoma wildtype (pLKO) or FADD knock down (shFADD) cells were treated with AS1842856. Cells were next collected, lysed and total protein probed for p21 protein expression. CCHOSD and HOS cells were treated with AS1842856 for 24 hrs. LM7 cells were treated with AS1842856 for 48hrs. **A**, Lentiviral-mediated knockdown of FADD. **B**, FADD knockdown alters p21 expression in OS cells. FADD knockdown reduces FOXO1 inhibition-induced p21 expression. **C**, CCHOSD cells, **D**, HOS cells, **E**, LM7 cells. Beta-actin served as a protein loading control. Immunoblot is representative of immunoblots from two independent experiments.

## 3. Discussion

Several studies have reported a correlation between cancer cell resistance to anticancer drug treatment and FOXO1 expression levels. For example, increased FOXO1 levels are associated with ovarian cancer cell resistance to paclitaxel [20], while ovarian cancer with down-regulated FOXO1 exhibits resistance to cisplatin [21]. These observations underscore the need to better understand the role of FOXO1 in the sensitivity or resistance of cancer cells to anticancer drug treatment. In the present study, Western blot analysis revealed that the OS cell lines studied express different basal levels of FOXO1 (Figure 1). Basal levels of FOXO1 showed a correlation between FOXO1 protein expression and sensitivity to CPT-induced cytotoxicity. Specifically, higher protein expression of FOXO1 correlated with increased sensitivity to CPT-induced cytotoxicity as indicated by the CPT doses used to induce cytotoxicity (Figures 1,3). This correlation was not observed in the two pancreatic cancer cell lines studied. Specifically, the basal level of FOXO1 was higher in AsPc1 cells compared to MiaPaCa2 cells; however, AsPc1 exhibited a slight decrease in sensitivity to CPT while MiaPaCa2 exhibited a slight increase in sensitivity to CPT (Figures 1,3,6). Although the decrease or increase in sensitivity was not significant, these results suggest that FOXO1 inhibition has a different effect on sensitivity to anticancer drug treatment within pancreatic cancer. This observation is significant because it suggests that FOXO1 expression may not be a reliable predictive indicator of the response of pancreatic cancer cells to anticancer drug treatment, namely CPT.

To investigate the effect that FOXO1 inhibition has on the response of OS to anticancer drug treatment, OS cells were pretreated with AS1842856 followed by CPT treatment. FOXO1 inhibition reversed CPT-induced cytotoxicity in all OS cell lines investigated (Figure 4), indicating that FOXO1 contributes to cell death following CPT treatment. Inhibition of FOXO1 also increased p21 expression and induced cell cycle arrest in all OS cells studied (Figure 2, 7). CPT induces cell death by causing DNA damage during DNA replication; therefore, it is plausible that the reversal in CPT-induced cytotoxicity mediated by FOXO1 inhibition was due to reduced cell cycle progression or cell cycle arrest facilitated by increased p21 expression.

The association of FOXO1 and FADD has been previously reported [16,17]. However, the understanding of this interplay on cell cycle regulation or response to anticancer drug treatment has not been investigated in OS. Before investigating the combined effect of FOXO1 inhibition and FADD knockdown on CPT-induced cytotoxicity, the effect of FADD knockdown alone on CPT-induced cytotoxicity was investigated. FADD knockdown increased OS sensitivity to CPT-induced cytotoxicity, suggesting that FADD serves a protective role following CPT treatment (Figure 5A). FADD status has been reported to have opposing effects on the response of cancer cells to anticancer drug treatment. For example, FADD knock down sensitizes pancreatic cancer to Adriamycin-induced cell death [10]. Conversely, FADD inhibition prevents cisplatin-induced cell death in HT29 and HTCT116 cells and prevents etoposide-induced apoptosis in U937 [22]. In addition, Jurkat cells with FADD knockout are resistant to doxorubicin or etoposide-induced apoptosis [23]. A comprehensive study that investigates the effect that FADD inhibition has on the response of different cancers to treatment with different anticancer drug classes will contribute to selecting the best anticancer drugs to use in combination therapy that includes FADD inhibition. FOXO1 inhibition significantly reduced the level of FADD knockdown-induced cytotoxicity (Figure 5B). This result indicates that FADD knockdown does not alter the tumor suppressor effect of FOXO1.

FOXO1 regulates the cell cycle in multiple cancers by regulating the expression of p16, p21 or p27 [24, 25]. To investigate the molecular mechanism responsible for FOXO1 inhibition-induced cell cycle arrest, basal level of p21 was determined and revealed a difference in the expression of p21 among the OS cell lines investigated. p21 is induced by DNA damaging agents and is a major CDKI that regulates cell cycle in cancer cells (26,27). Indeed, CPT treatment increased p21 expression in all OS cell lines investigated (Figure 3). FOXO1 is well documented to be a positive regulator of p21 expression [24]. Therefore, it was expected that FOXO1 inhibition would decrease p21 expression. However, in the present study, FOXO1 inhibition increased p21 expression in all OS cells (Figure 2). It is plausible that the FOXO1 inhibition-induced increase in p21 was responsible for the observed cell cycle arrest and reversal in CPT-induced cytotoxicity. The observed FOXO1 inhibition-induced increase in p21 expression was reversed by FADD knock down (Figure 9C-9E). The observation that FADD knock down reversed FOXO1 inhibition-induced increase in p21 expression suggests that FOXO1 and FADD have opposing effects on p21 expression.

## 4. Materials and Methods

### 4.1. Cell Lines, Cell Culture and Reagents

HOS is a non-metastatic human OS cell line. CCHOSD and LM7 are metastatic human OS cell lines. AsPc1 and MiaPaCa2 are metastatic human pancreatic cell lines. Cells were treated with AS1842856 to inhibit FOXO1 transcriptional activity. Lentiviral shRNA targeted against FADD RNA was used to knock down FADD protein expression in the OS cells. Lentivirus containing empty shRNA vector served as the control for FADD knockdown OS cells. A detailed description of lentivirus generation was provided previously [28]. Cells were cultured in a CO_2_ incubator maintained at 37°C and 5% humidity. Cells were treated with drug as indicated in figures and figure legends. AS1842856 was purchased from (Millipore Sigma, Burlington, MA). Camptothecin (CPT) was purchased from ChemWerth (Woodbridge, CN). Beta-actin antibody was purchased from Sigma-Aldrich (St. Louis, MO). p21 antibody was purchased from Cell Signaling Technology (Danvers, MA). GAPDH antibodies were purchased from Santa Cruz Biotechnology (Dallas, TX). Fetal bovine serum (FBS) and Dulbecco’s Modified Eagle Medium (DMEM) were purchased from VWR International (Radnor, PA). Cell culture supplements were purchased from Invitrogen (Carlsbad, CA).

### 4.2. Cytotoxicity and cell cycle analysis

Cytotoxicity was determined by assessment of the cell population in the sub G0/G1 area of the cell cycle which is considered non-viable [29]. Propidium iodide (PI) is a dye that binds to double stranded DNA by intercalating between base pairs. Cells with degraded DNA (non-viable) have insufficient PI binding and appear in the subG0/G1 population of the cell cycle histogram. Propidium iodide binding to DNA also allows for assessment of cell cycle status [29]. Cells in G0/G1 have DNA that is 2N and have less PI staining than cells in S stage that are between 2N and 4N. Cells in G2/M stage have completed DNA synthesis and are 4N; thus, having the greatest PI staining and farthest shift to the right in the histogram. Following drug treatment, supernatant and cells were collected and centrifuged at 500 g for 5 min at 4°C. The resultant pellet was fixed with cold 70% ethanol and incubated for 24 h at -20°C. Following fixation, the cells were washed with PBS and centrifuged to remove the ethanol and next incubated with PI for 24 h followed by analysis on a flow cytometer.

### 4.3. Western Blot Analysis

Following drug treatment, supernatant and cells were collected and centrifuged at 500 g for 5 min at 4°C. The resultant pellet was lysed with RIPA lysis buffer containing protease and phosphatase inhibitor cocktail and centrifuged at 14,000 g for 15 min at 4°C. Supernatants were then collected, and total protein was determined by BioRad reagent (BioRad Laboratories, Hercules, CA). 15ug - 30ug of protein were resolved in SDS-polyacrylamide gels (SDS-PAGE) and transferred onto nitrocellulose membranes (BioRad Laboratories, Hercules, CA). Membranes were blocked with 5% nonfat milk and next incubated with primary antibody. Membranes were next washed and incubated with appropriate secondary antibody conjugated to HRP. Following secondary antibody incubation, membranes were washed, and signal detected with ECL detection reagent (Santa Cruz Biotechnology, Inc, Dallas, TX). Beta-actin or GAPDH served as protein loading control.

### 4.4. Statistical Analysis

Results are presented as means + standard error mean (SEM). Experimental data were analyzed using 2-tailed Student *t* test. P values less than 0.05 were considered statistically significant.

## 5. Conclusions

The results of this study indicate that FOXO1 has a pro-death function in OS treated with CPT. The results also indicate that the FOXO1 inhibition-induced reversal of CPT-induced cytotoxicity is cancer type-dependent. These conclusions are based on the observation that inhibition of FOXO1 significantly reversed CPT-induced cytotoxicity in all OS cell lines investigated but not in the pancreatic cancer cell lines. The results also suggest that FADD counters the effect of FOXO1 on p21 protein expression in OS. This conclusion is based on the observation that FADD knockdown reduced FOXO1 inhibition-induced increase in p21 expression. To the best of our knowledge, this is the first report of AS1842856-mediated inhibition of FOXO1 reversing CPT-induced cytotoxicity in OS. This is also the first report of FOXO1 inhibition or FADD knock down altering the expression of p21 in OS. The results of this study underscore the need to investigate further the individual and combined role of FOXO1 and FADD in cell cycle regulation and cancer cell resistance or sensitivity to anticancer drug treatment.

## Author Contributions

Conceptualization, M.G.H.; Methodology, D.W., A.H., A.B., M.G.H.; Writing – original draft preparation, D.W., M.G.H.; literature check, manuscript review and editing, D.W., N.G., D.R.; Supervision, M.G.H; Funding acquisition, M.G.H. All authors have read and agreed to the published version of the manuscript.

## Data Availability Statement

The data that support the findings of this study are available from the corresponding author upon reasonable request.

## Abbreviations

The following abbreviations are used in this manuscript:

FOXO: forkhead box-O
FADD: fas-associated death domain
CDK: cyclin-dependent kinase
CDKI: cyclin-dependent kinase inhibitor
CPT: camptothecin

